# Uncovering Astrocyte Morphological Dynamics Using Optical Diffraction Tomography and Shape-based Trajectory Inference

**DOI:** 10.1101/2024.08.16.608356

**Authors:** Pooja Anantha, Piyush Raj, Emanuela Saracino, Joo Ho Kim, Jeong Hee Kim, Annalisa Convertino, Luo Gu, Ishan Barman

## Abstract

Astrocytes, integral components of the central nervous system, are increasingly recognized for their multifaceted roles beyond mere support cells. Despite their acknowledged importance, understanding the intricacies of astrocyte morphology and dynamics remains limited. Our study marks the first exploration of astrocytes using optical diffraction tomography (ODT), aiming to establish a solid foundation for their detailed characterization. It offers valuable insights into the morphological changes in postnatal rat cortical astrocytes over a 7-day *in vitro* period in a label-free manner. Through comprehensive analysis of 3D refractive index maps and shape characterization techniques, we elucidate the developmental trajectory and dynamic morphological transformations of astrocytes in culture. Specifically, our observations revealed increased area and transition to larger, flattened shapes, and alterations in cell volume and density, indicating shifts in cellular composition. Furthermore, by employing unsupervised clustering and pseudotime trajectory analysis, we tracked the morphological evolution of astrocytes from highly directional to evenly spread shapes. These results emphasize the dynamic nature of astrocytes. In addition, this analysis marks the first use of trajectory inference based solely on morphology for neural cell types. Future studies could employ ODT to examine the morphological dynamics and interactions of various neural cell types on other types of substrates.

## 1. Introduction

Astrocytes, a fundamental type of glial cell within the central nervous system (CNS), hold a pivotal position in the intricate tapestry of brain and spinal cord function. In recent years, the spotlight on astrocytes has intensified, revealing their pivotal roles in information processing within the CNS, as proposed by the tripartite synapse hypothesis [1], [2], [3], [4], [5], [6]. This heightened focus stems from a growing acknowledgment of astrocytes as more than mere support cells; they are active participants in the neural dialogue, integral to the formation and functioning of synapses and neural networks. Indeed, they play a major role in providing structural scaffolding, facilitate the maintenance of neurotransmitter balance and ionic concentrations in the synaptic cleft, regulate cerebral blood flow, and actively contribute to the formation of the blood-brain barrier (BBB). Their multifaceted responsibilities extend to synaptic modulation, neuroprotection, metabolic support to neurons, and participation in the immune response within the CNS [7], [8], [9], [10], [11], [12]. Perturbations or pathological alterations in astrocyte activity have been associated with a spectrum of neurological disorders, encompassing Alzheimer’s disease, Parkinson’s disease, epilepsy, and multiple sclerosis [10], [11], [13]. Hence, a thorough comprehension of astrocytes and their multifaceted contributions is highly desired with significant implications for the development of novel therapies to treat neurological disorders [10], [11], [12], [13].

However, despite their recognized importance, our understanding of astrocytes remains limited with a particularly intriguing feature being their extraordinary morphological complexity. Generally, they are considered as “star like”-shape cells, while specifically in gray matter, these cells extend numerous fine, densely branching processes, allowing intimate interaction with neurons and synapses, enabling precise regulation of synaptic activity and neurotransmitter levels. Conversely, in white matter, astrocytes possess fewer and less complex processes, yet their role in maintaining structural integrity and supporting axonal function remains critical. This morphological intricacy is a window into the vast spectrum of astrocytic functions, from synaptic regulation to structural support [12], [14], [15], [16], [17]. All these tasks necessitate close and highly adaptable contacts between astrocytes and their targets, interactions that can dynamically change based on individual physiological conditions. Yet very little is known about the development of the stunning complexity of astrocytes from the thin precursor glial cells and the molecular machinery behind these shapeshifts.

Central to the significance of understanding astrocytes is their volume – a key determinant of their structural support function. Their volume is linked to their structural support function, primarily through numerous fine processes that extend and interact with neurons, synapses, and blood vessels [18], [19]. Furthermore, the ability of brain cells to regulate their volume is mediated essentially by astrocytes’ homeostatic function. Constantly, water and ion exchanges occur in the extra and intra-cellular milieu during the brain network activity and the regulation of the brain hydrosaline composition is tightly dependent by the characteristic star-like morphology and protein expression of *in vivo* astrocytes. They are equipped with ion channels and transporters that guide both the water flux the organic osmolytes and salts across the plasma membrane. Moreover, astrocyte volume function governs their ability to regulate neurotransmitters like glutamate effectively, profoundly impacting synaptic transmission and overall neuronal function [20], [21], [22], [23], [24]. The intricate dance of ionic balance and homeostasis within the CNS, critical for optimal neuronal function, is also influenced by astrocyte volume. Notably, changes in astrocyte volume can be indicative of trauma, inflammation, or neurodegenerative diseases, offering valuable diagnostic insights into the underlying pathological processes [25], [26].

Several research studies highlighted the significance of investigating astrocyte volume regulation mechanism both in case of cell swelling and cell shrinkage. Upon the following mechanism has been defined as Regulatory Volume Decrease (RVD) and Increase (RVI). Among water and ion channels the aquaporin-4 (AQP-4) and Volume Regulated Ion Channels (VRAC), expressed by astrocytes, has been shown as critical and crucial in *in vivo* recovery of physiological cell volume during and after a brain volume alteration. VRAC control the efflux of Chloride (Cl-) and organic osmolytes (such as taurine, glutamate, and aspartate) [27]. Recent studies have identified the protein called LRRC8-A in the plasma membrane of primary cortical astrocytes and in situ at the perivascular interface with endothelial cells as key factor for astroglial volume homeostasis. The VRAC and AQP4 functional interaction has been demonstrated in primary astrocytes. Notably, the alteration in expression and function of AQP4, VRAC and calcium channels such as Transient Receptor Potential Vanilloid 4 Channels (TRPV4) is widely recognized as pathogenic in neurological conditions characterized by dysregulation of astrocytic homeostasis [28], [29].

Quantifying astrocyte volume accurately remains a challenge and existing studies primarily rely on fluorescence, which does not provide precise quantitative metrics and may potentially lead to inaccurate volume calculations [30], [31], [32]. Furthermore, exploring astrocyte mass has been overlooked, primarily because there are relatively few methods that can accurately measure single-cell mass over time in living cells. Although cell mass and volume are generally tightly correlated, it is known that volume change can be transient and decoupled from cell mass at different stages of differentiation or of the cell cycle [33], [34], [35], [36]. Because astrocytes directly impact the organization and stability of neuronal networks, cell mass may provide crucial insights during their evolution from precursor glial cells to mature astrocytes. Additionally, a change in astrocyte mass could reflect alterations in metabolic support, potentially impacting neuronal energy metabolism and overall brain function. Thus, delving into the quantification of astrocyte volume and mass becomes an essential endeavor, paving the way for a deeper comprehension of their varied roles in CNS health and pathology.

In this pursuit, recent advancements in optical methods, particularly holographic microscopy, are emerging as potentially transformative methodologies. Holographic microscopy, encompassing quantitative phase imaging (QPI) in a broader sense [37], [38], [39], offers a means to gain precise morphological and biochemical insights without the need for external labeling. QPI involves the measurement of optical field images, capturing nanoscale distortions of wavefronts passing through a sample using laser-based interferometry. Beyond the amplitude images provided by conventional intensity-based microscopy, QPI allows for the quantitative measurement of optical phase delay maps, intimately tied to the refractive index (RI) distribution of the sample. A variant of QPI, optical diffraction tomography (ODT), facilitates the acquisition of 3D refractive index (RI) maps of cells [40], [41], [42], [43]. Crucially, QPI also allows accurate mass measurements of adherent cells down to a precision of less than 10 picograms [44].

Leveraging the quantitative ODT signals based on intrinsic contrast, our current study seeks to address critical gaps in visualizing and analyzing astrocyte morphology. By non-destructive imaging of the individual unstained cells, we assess astrocyte volume as well as cell mass in postnatal rat cortical astrocytes over the course of a week (**Fig. 1**). Employing both 3D and 2D RI maps obtained, this investigation marks the first use of ODT for studying astrocytes, to the best of our knowledge. Additionally, our research presents a new method for understanding the developmental progression and morphological transformations of astrocytes in culture over time, employing the Zernike polynomial method for shape characterization to unravel the developmental trajectory. The immense potential of ODT showcased in this study lays a strong foundation for future investigations, offering a pathway to unlock deeper insights into astrocyte biology and its implications in the realm of neuroscience and neurological disorders.

**Figure 1:**
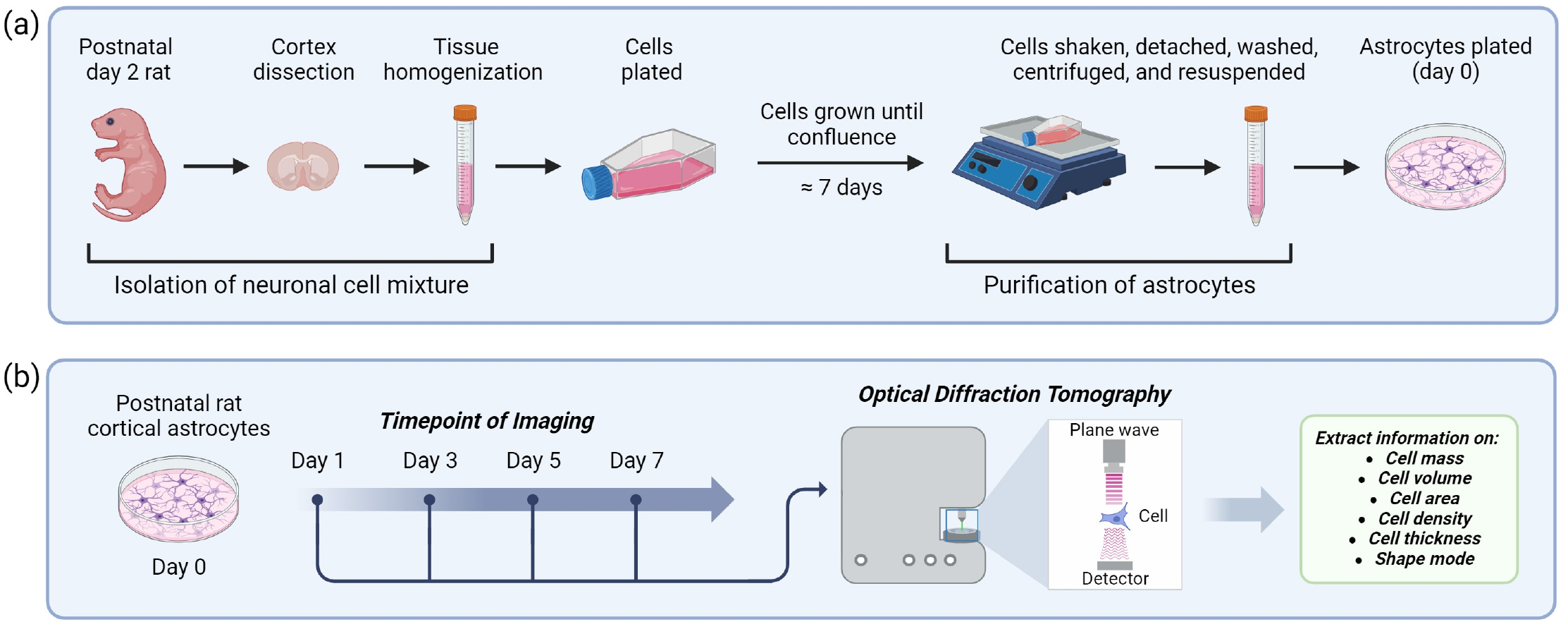
Project overview. a) Astrocytes were extracted from postnatal day 2 rat cortices by initially isolating the neural cell mixture followed by purification of astrocytes. B) ODT was utilized to study the heterogeneity and quantify morphological changes of astrocytes for 7 days.

## 2. Methods

### 2.1. Preparation and purification of postnatal rat cortical astrocytes

All animal experimental procedures were conducted in strict accordance with Animals Care and Use Committee at Johns Hopkins University. Astrocytes were extracted from postnatal day 2 Sprague-Dawley rats by initially isolating neural cell mixture from the rat followed by purification for astrocyte extraction [45], [46]. The meningeal layer was removed, and the cortex was dissected in cold HBSS (Thermofisher) with 2% P/S. The wet cortical tissue was then placed in a petri dish for chopping with a sterile blade. The chopped tissue (∼1 mm^3^ pieces) was transferred to pre-warmed 0.05% trypsin-EDTA (Thermofisher) and incubated at 37°C shaken every 5 minutes. After 10 minutes, trypsin was quenched using 10% heat-inactivated fetal bovine serum (Cytiva), 100U/ml penicillin, and 100U/ml streptomycin (Thermofisher), and Dulbecco’s modified Eagle’s medium with glucose and pyruvate (DMEM, Thermofisher) solution (cDMEM). The tissue was triturated, centrifuged, and the pellet was resuspended in cDMEM. The cell suspension was further homogenized, strained through a cell strainer, and the process was repeated total of three times. Mixed neural cells were seeded onto poly-D-lysine (PDL, Thermofisher)-coated T75 culture flasks (one brain per flask). PDL coating involved incubating flasks with 10μg/mL PDL, diluted in deionized (DI) water, and added to each dish at a volume of 0.125mL/cm^2^ for 30 minutes at 37°C, followed by two washes with DI water for 5 minutes each. Initial seeding required 15 ml of culture medium, followed by 24 hours of culturing with subsequent 100% medium exchange to remove excess cell debris. The culture was maintained until cells reached over 90% confluence, which usually required a week, with media changes every 3 days. When cells were ready for purification, the flask was sealed, covered, and shaken on orbital shaker at 210 rpm for 1 hour in 37°C incubator then vigorously shaken around 10 times to dislodge remaining microglia and oligodendrocytes. The shaking focused on the flask’s bottom to ensure detachment. Following shaking, media was aspirated, monolayers were washed with DPBS (Thermofisher), and microscopy confirmed non-astrocyte cell removal. Trypsin treatment, neutralization, and detachment were performed, and the cell suspension was transferred to tubes. Additional washes, centrifugation, and resuspension in serum-free DMEM were carried out. Cell counting utilized a trypan blue solution on a Countess slide. Astrocytes were then re-plated on the 1.5H glass bottom petri dishes. The day of replating is considered day 0. Once seeded onto dishes, astrocytes were cultured and fixed at 4 different time points, ranging from day 1 to day 7. Fixation involved 15 minutes of incubation at room temperature with 4% paraformaldehyde (PFA, Biotium) followed by three washes with DPBS for 5 minutes each. Three dishes, or three repeats, were fixed per “day”, for a total of 12 dishes.

### 2.2. Immunofluorescence staining and astrocyte purity assessment

The characterization of astrocytes involved immunostaining for glial fibrillary acidic protein (GFAP) and imaging via QPI. The staining process allowed for the visualization and identification of astrocytes based on their expression of GFAP. The astrocytes were fixed with 4% PFA for 20 minutes at room temperature, followed by three washes with DPBS for 5 minutes each. Fixed cells were then permeabilized with 0.2% Triton X-100 (Sigma-Aldrich) for 30 minutes at room temperature, and then blocked with blocking buffer composed of DPBS with 5% normal goat serum (Thermofisher), 0.2% Triton X-100, and 1% bovine serum albumin (BSA, Sigma-Aldrich) for 1 hour at room temperature. Then, cells were incubated with GFAP (1:1000, Dako), diluted in blocking buffer overnight at 4°C. After the incubation, cells were washed with DPBS for 10 minutes three times and incubated for an hour with the secondary antibody, Alexa Fluor 488 (1:1000, Thermofisher), diluted in the blocking buffer at 4°C. Astrocytes were also stained with Hoechst 33342 (1:2000, Thermofisher) and washed three times with DPBS for 5 minutes each and kept in dark until imaging.

### 2.3. Immunofluorescence staining of aquaporin-4 (AQP-4) and volume-regulated anion channel (VRAC)

Astrocytes were plated onto six 1.5H glass bottom petri dishes. Three dishes were fixed on day 1 and the other three dishes were fixed on day 7. Fixation involved 20 minutes of incubation at room temperature with 4% PFA followed by three washes with DPBS for 5 minutes each. Following fixation and washing, cells were incubated in blocking buffer composed of DPBS with 0.2% Triton X-100 and 1% BSA for 30 minutes at room temperature. Next, they were incubated in the primary antibody solution composed of anti-aquaporin 4 (AQP-4) (1:300 dilution in blocking buffer, Millipore Sigma) and anti-LRRC8A antibody (1:200 dilution in blocking buffer, Alomone) for 3 hours at room temperature followed by three washes with DPBS for 5 minutes each. Then, they were incubated in the secondary antibody solution composed of Alexa Fluor 568 (1:1000, Thermofisher) and Alexa Fluor 488 (1:1000, Thermofisher) followed by three washes with DPBS for 5 minutes each. Lastly, astrocytes were stained with Hoechst 33342 (1:2000, Thermofisher) and washed three times with DPBS for 5 minutes each and kept in dark until imaging.

### 2.4. Optical diffraction tomographic imaging and analysis

ODT was performed to capture astrocyte morphology on 1, 3, 5, and 7 days (DIV) after seeding. After fixation, astrocytes were captured using a holotomography system (HT-2, TomoCube, Republic of Korea) comprised of a motorized stage, a 60x, 1.2 NA water-immersion objective, a 532 nm laser, and a dynamic micromirror device, that measure the 3D refractive indexes. TomoStudio (TomoCube, Republic of Korea) was used to visualize and obtain 3D RI tomogram images and 2D maximum intensity projection (MIP) images. TomoStitcher (TomoCube, Republic of Korea) was used to stitch the images. Image processing was done using ImageJ. Cell segmentation was performed manually on 2D MIP images using ImageJ. After cell segmentation, we obtained about 450 astrocytes per timepoint. These 2D segmented cells served as masks and were applied to the stacks of 3D RI tomogram images to generate 3D RI tomograms featuring only one cell per field of view [47]. Typically, the original 3D RI tomogram images contain multiple cells. Quantitative analyses were performed on 3D RI tomogram images with one cell per field of view using MATLAB. Area was calculated by counting the number of nonzero pixels in the masks. Cell dry mass values were obtained by initially setting refractive index thresholds and then calculating the surface integral of the optical path difference over the specific refractive index increment [42]. Volume was calculated by counting the number of nonzero pixels in 3D segmented images.

For cellular trajectory inference, we used the protocol developed in our recent work [48]. Briefly, we used Zernike polynomials in two-dimension to convert 2D shape to one-dimensional vector. The 2D image was centered and aligned along its major axis. The image was then converted in a polar coordinate system (r, θ) from cartesian system. After that, we used Zernfun [49] to compute the Zernike functions Znm (r, θ) which gave us the value of Zernike moments. We used trajectory inference (TI) method to perform pseudo-temporal ordering of different shapes. Scfates [50] and Scanpy [51] toolbox was used for this analysis.

## 3. Results and discussion

### 3.1. Postnatal rat cortical astrocytes undergo dynamic morphological changes over a period of 7 DIV

In this comprehensive 7-day investigation, the astrocytes showcased a remarkable spectrum of changes not only in their morphological structure but also in crucial physical parameters such as mass, volume, density, area, and thickness. These dynamic alterations offer profound insights into the adaptability and dynamic behavior of astrocytes within an *in vitro* environment.

Illustrated in Figure 2, the optical diffraction tomography (ODT) images of postnatal day 2 rat cortical astrocytes provide a visual narrative of these alterations. The astrocytes were initially subjected to immunostaining for glial fibrillary acidic protein (GFAP), a commonly used marker for astrocytes (**Fig. 2a**). This confirmed that over 95% of the cells in the sample were indeed astrocytes. The subsequent ODT measurements carried out on a different set of astrocytes at various time points—DIV 1, 3, 5, and 7—reveal a fascinating trajectory of changes in astrocytic physical parameters (**Fig. 2b**). The quantification of these parameters is presented in **Figures 3a-c**.

**Figure 2:**
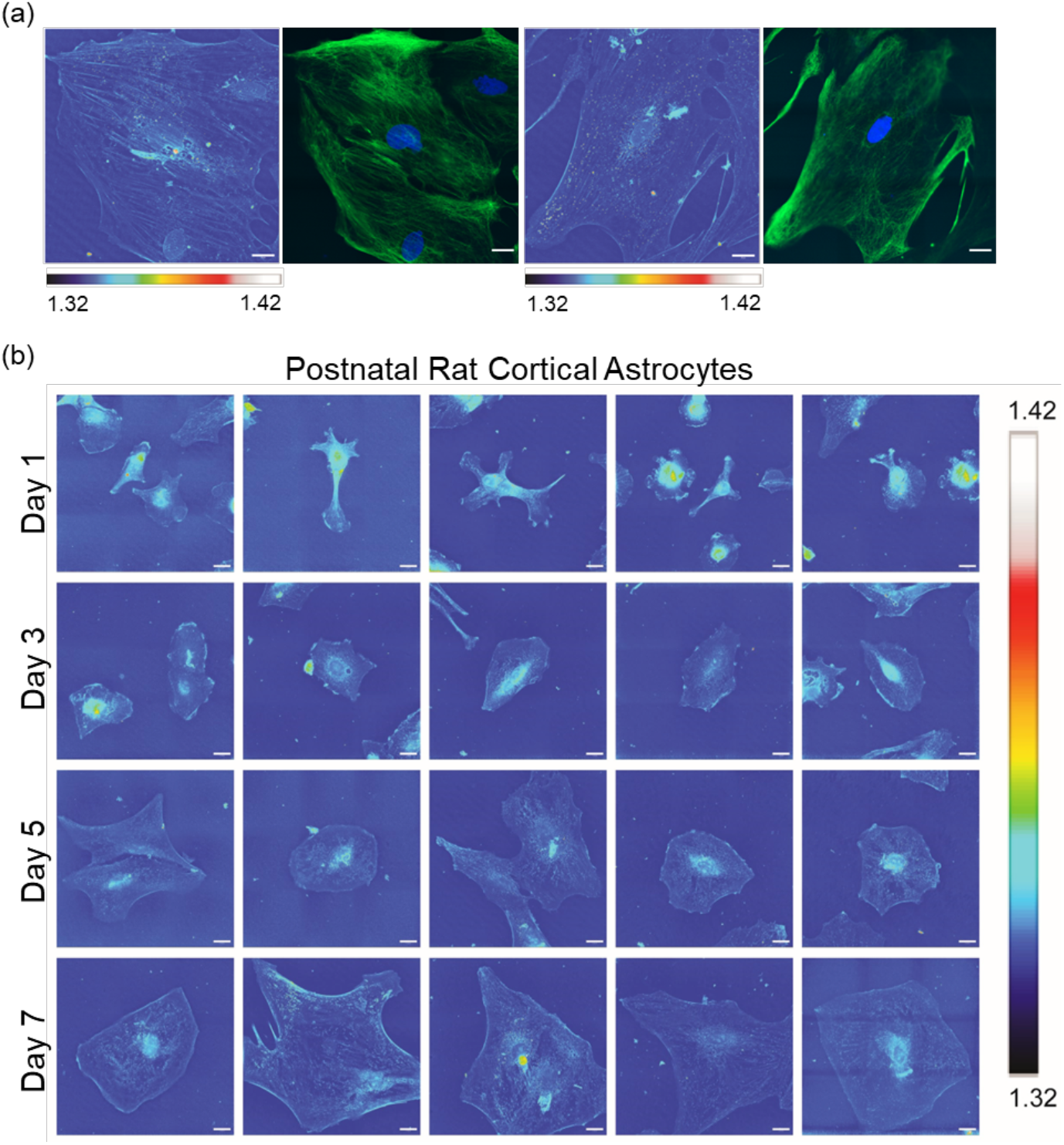
ODT images of postnatal day 2 rat cortical astrocytes a) stained for GFAP (green) and nucleus (blue), (left side image is the 2D MIP representation) and b) at various time points (scale bar: 20 μm). Legend indicates RI value range.

**Figure 3:**
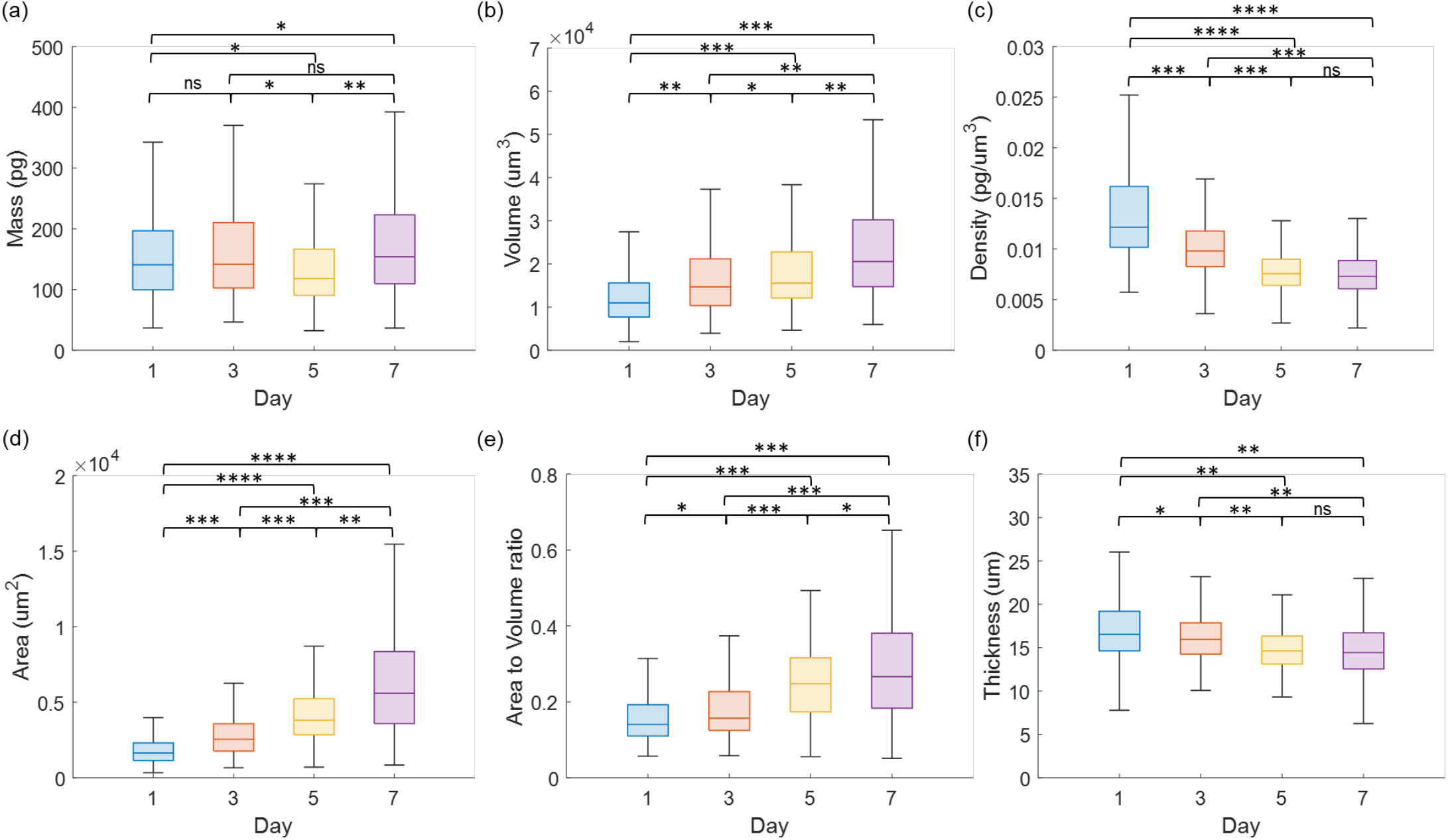
Quantification of physical parameters of postnatal day 2 rat cortical astrocytes (*p ≤ 0.01 and effect size ≤ 0.2, **p ≤ 0.001 and 0.2 < effect size ≤ 0.4, ***p ≤ 0.0001 and 0.4 < effect size ≤ 0.7, ****p ≤ 0.00001 and effect size > 0.7, not significant (ns): p>0.01). Complete list of p-values and effect size values are provided in SI.

Primarily, a notable observation was the progressive increase in the area covered by astrocytes. Commencing from the early stages (1 DIV) with a radius spanning approximately 50 to 100 μm, the astrocytes exhibited a gradual expansion in their coverage, eventually displaying a planar and flattened morphology by days 5 and 7. 3D orbital views of a 1 DIV astrocyte and a 7 DIV astrocyte are provided in SI (**Video S1** and **Video S2**). This observed expansion indicates cell growth and spreading, influenced by the composition of the culture medium and the properties of the substrate. The broader expanse covered by astrocytes underscores their ability to adapt and dynamically interact with their surrounding milieu—a fundamental aspect pivotal to their role in supporting and modulating neural activities.

### 3.2. Precise measurement of astrocyte morphology reveals an substantial increase in volume and cell area with no concurrent changes in dry mass

Grasping the intricacies of shifts in astrocyte mass, volume, and area holds profound importance in comprehending the dynamic behavior and adaptability of astrocytes within a controlled *in vitro* milieu. These transformations not only underscore the influence of external factors such as culture conditions but also illuminate the nuanced cellular processes and mechanisms that drive astrocytic behavior. The observed alterations provide a glimpse into the extraordinary plasticity of astrocytes, hinting at their capacity to dynamically respond to variations in their environment. Moreover, delving into the specific biochemical and biophysical changes that accompany these alterations in physical parameters holds promise for unlocking deeper insights into the functional roles of astrocytes in the nervous system. The intricate interplay between structural modifications and functional adaptations in astrocytes remains an area ripe for further exploration. Quantification of astrocyte morphological parameters are shown in **Figure 3**.

As depicted in **Figure 3a**, there is no substantial alteration in the dry mass of astrocytes over the 7-day period (effect size < 0.4). However, there is a discernible increase in volume (**Fig. 3b**) coupled with a decrease in density (**Fig. 3c**), indicating the uptake of non-dry components. This process contributes substantially to the observed increase in cell area (**Fig. 3d**) from day 1 to day 7. The flattening of astrocytes is also exemplified by a slight reduction in thickness (**Fig. 3f**). In addition, it is important to note that the thickness experiences only a minor decrease, while volume is concurrently increasing. Given that volume is a product of area and thickness, it implies that the rapid expansion in area effectively overcompensates for the slight reduction in thickness (**Fig. 3e**). The quantification of astrocyte morphological parameters with the precision demonstrated in our study represents a major contribution to existing research. To our knowledge, no prior studies have ventured into such detailed measurements of astrocyte dry mass, volume, density, area, and thickness.

In our pursuit of a deeper understanding surrounding the increase in volume without a substantial increase in mass within astrocytes over the 7-day interval – we focused our scrutiny on specific channels, namely aquaporin-4 (AQP-4) and volume-regulated anion channel (VRAC), which have been known to play a major role in astrocyte volume regulation [31], [32], [52]. *In vivo* astrocytes act by both these transmembrane proteins at the nano-microdomains of their end-feet and by calcium signaling for the synaptic plasticity, memory, learning, and regulation of sleep and metabolism. Specifically, AQP-4 is consistently found in high concentrations in the plasma membrane of perivascular glial processes in astrocytes *in vivo*, and its expression undergoes changes in specific pathological conditions linked to brain edema or altered glial migration. When astrocytes are cultured, they deviate from their characteristic star-like shape and the continuous plasma membrane localization of AQP4 observed *in vivo* [27]. Equally, we address to investigate on VRAC expression in astrocytes by performing immunofluorescence experiments on AQP-4 and VRAC. Recent studies in cultured astrocytes have found that LRRC8A, a leucine-rich-repeat subunit is a main constituent of the VRAC transport complex [31], [32], [53]. The following unit is indispensable for the swelling-induced and ATP-induced release of neuroactive molecules. The astrocytic VRAC results in several physiological processes and human pathologies given its critical role in cell volume regulation [54]. In this view, we performed immunostaining for AQP-4 and LRRC8A at 1 and 7 DIV astrocytes and compare the data to show a possible different expression of both the water and ion proteins as responsible of the volume difference observed in **Figure 3e**. However, as depicted in **Figure S1**, the expression of both AQP-4 and VRAC are found throughout the cell but appears to be concentrated more around the nucleus, with some presence observed at the cell boundary for both 1 and 7 DIV astrocytes. Both comparison of AQP-4 and VRAC stained astrocytes reveals that no significant difference can be observed in the cell proteins expression at 1 and 7 DIV. However, given the prominent role of AQP4 in the water permeability and cell volume regulation in astrocytes to get further insights we observed that a higher but not significant increase of AQP-4 (red staining) is expressed in the boundary of astrocytes at 7 DIV compared to 1 DIV. In agreement with previous works, the expression of AQP4 in polygonal astrocytes *in vitro* is mislocalized [55].

Despite the previously discussed marked increase in cell area, the concomitant changes in AQP-4 and VRAC expression relative to cell area remain inconclusive. This experimental evidence suggests the existence of additional contributing factors that induce alterations in volume without a corresponding impact on the cellular dry mass. Although our current study does not delve into these factors, addressing them may be a promising avenue for future investigations.

### 3.3. Further morphological analyses using Zernike polynomials provide novel insights

Cellular trajectory inference was performed from morphological parameters using Zernike polynomial. In the past, cellular morphological analysis primarily relied on classification algorithms or the categorization of cells into distinct shape modes in order to differentiate them based on their pathophysiological conditions [43], [56], [57], [58], [59]. However, numerous biological processes, including cell differentiation and development involve a continuous and asynchronous transition metaphorically depicted by Waddington’s landscape plot [60], [61], [62], [63]. To capture the intricacies of the development process through morphological analysis, we converted the cell mask to equivalent Zernike moment. We selected the method of using Zernike polynomial for its ability to faithfully reconstruct cell shapes while maintaining rotational invariance. Thereafter, we use trajectory inference toolbox (TI) which has originally been developed for single cell transcriptomics studies. This is a first of its kind study for any neuronal cells to track cellular lineage through morphology alone.

As previously discussed, the discernible contrast in astrocyte morphology between day 1 and day 7 is evident; however, the morphological alterations between consecutive days appear subtle. This observation is effectively captured in the UMAP-based supervised label plot (**Fig. 4a**), where a distinct separation between different days proves challenging. This observation aligns with our previous findings, illustrating that individual cells exhibit varied responses to the differentiation process, with some cells responding robustly while others respond more moderately to the same stimulus.

**Figure 4:**
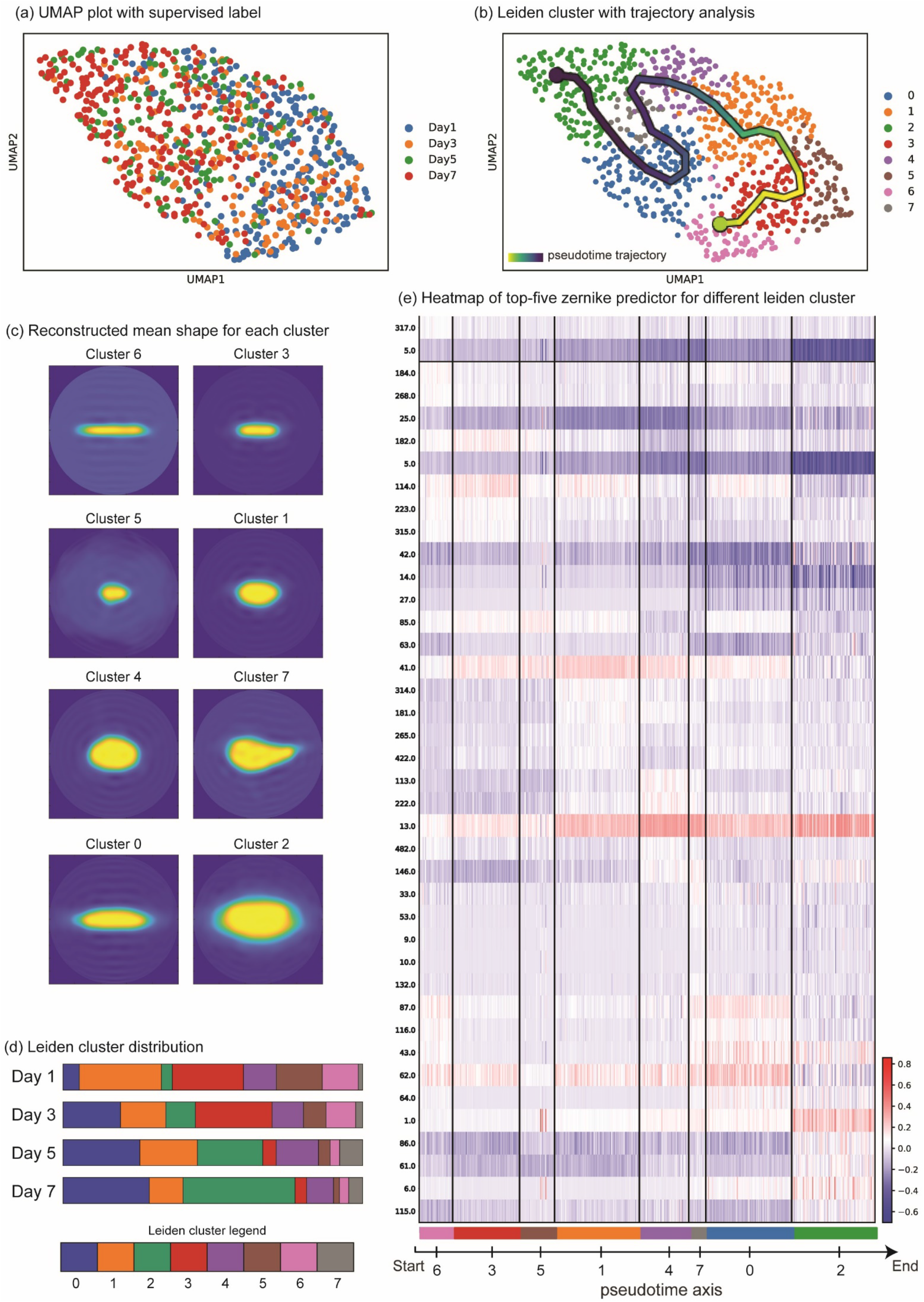
(a) UMAP plot from Zernike moment of cell mask images with supervised labels. (b) UMAP plot from Zernike moment of cell mask images with unsupervised leiden cluster labels. A pseudotime trajectory is also plotted on the curve. (c) Reconstructed mean shape of each identified clusters. (d) Leiden cluster distribution across different days. (e) Heatmap of top 5 variables from each cluster which were responsible for identification of clusters that are aligned on a pseudotime axis. The variable number on the left stands for Zernike polynomial index and each row corresponds to the value of Zernike moment for the given polynomial for all the images in the dataset. The solid black line along the y-axis is the margin for each cluster.

In **Figure 4b**, an unsupervised Leiden clustering was performed. Unconstrained by human labels, this clustering focuses solely on phenotypic responses, avoiding overfitting or an attempt to conform to supervised labels. This approach enables the identification of a continuum of phenotypes connected through pseudotime trajectory. The Leiden cluster plot, coupled with trajectory analysis, provides insights into how the structure evolves from one differentiation point to another (**Fig. 4b-d**). The average astrocyte shape initiates with a highly directional form in clusters 6 and 3 (**Fig. 4c**). As it progresses to cluster 1, the shape becomes more rounded and evenly spread. Day 1 astrocytes exhibit an almost equal distribution between cluster 3 and cluster 1, with a sparse population of other clusters, indicating the heterogeneity of the initial population. These clusters also appear ahead on the pseudotime trajectory (**Fig. 4e**). Progressing to day 3, day 5, and finally to day 7, there is a gradual increase in the percentage of cells from clusters 0 and 2, accompanied by a decrease in the percentage of clusters 3 and 1, which were predominant on day 1. In Figure 3a, the increase in the area of astrocytes is observed as the days progress; however, simple geometric parameters fail to capture the gradual shape changes illustrated in the reconstructed mean shape of each cluster (**Fig. 4c**).

Most astrocytes commence with a high aspect ratio shape and, in some cases, a small and rounded shape, transitioning to a more rounded morphology. The isotropic growth in 2D cell culture is attributed to the absence of constraints from the extracellular matrix for directional growth, as evident in cluster 2. A considerable number of astrocytes also retain the high aspect ratio shape, although they increase in area, as demonstrated in cluster 0. In **Figure 4e**, a heatmap of the top-five Zernike predictors for each identified cluster is plotted. Each pixel represents the value of the Zernike moment for a specific cell in that cluster. The gradual changes in the value of Zernike moment along the pseudotime axis are observed for several polynomials such as 5, 41, 13, and 1, among others. This analysis aids in identifying the variables responsible for differentiating the clusters from other shapes in a quantitative manner, offering valuable insights for building a concise model if a training model for morphology prediction is to be developed.

## 4. Conclusion

Our study represents an initial exploration into the realm of astrocyte examination through ODT, driven by the fundamental goal of creating a robust foundation for the characterization of astrocytes. It provides valuable insights into the morphological changes observed in postnatal rat cortical astrocytes over a 7-day period *in vitro*. Through detailed quantification using ODT, we observed notable alterations in astrocytic physical parameters, including mass, volume, density, area, and thickness. Importantly, while we noted a significant increase in cell area and volume, no concurrent changes in dry mass were observed, highlighting the complexity of cellular processes driving astrocytic behavior. Our findings underscore the adaptability and plasticity of astrocytes, essential for their roles in supporting neural activities. Crucially, ODT facilitates the extraction of quantitative data related to morphological parameters that were previously unexplored.

Moreover, our application of Zernike polynomials for morphological analysis reveals a continuum of phenotypic responses in astrocytes over time, emphasizing the dynamic nature of astrocytic differentiation. Through unsupervised clustering and pseudotime trajectory analysis, we delineate the evolution of astrocyte morphology from highly directional forms to more rounded, evenly spread shapes. These findings underscore the heterogeneity and dynamic nature of astrocytes. Moving forward, our study lays a strong foundation for future research endeavors aimed at deciphering the intricate interplay between structural modifications and functional adaptations in astrocytes, with profound implications for understanding CNS pathology. Future studies may involve the utilization of ODT to examine morphological dynamics and interactions of other neural cell types on various substrates.

## Acknowledgements

The authors acknowledge support from the Air Force Office of Scientific Research (FA9550-22-1-0334). Figure 1 was created with BioRender.com.

