## Supplementary for "Uncovering Astrocyte Morphological Dynamics Using Optical Diffraction Tomography and Shape-based Trajectory Inference": Astro_SI_PA_05162024.pdf

**Conflicts of Interest**

The authors declare no potential conflicts of interest.

### Supplemental Information

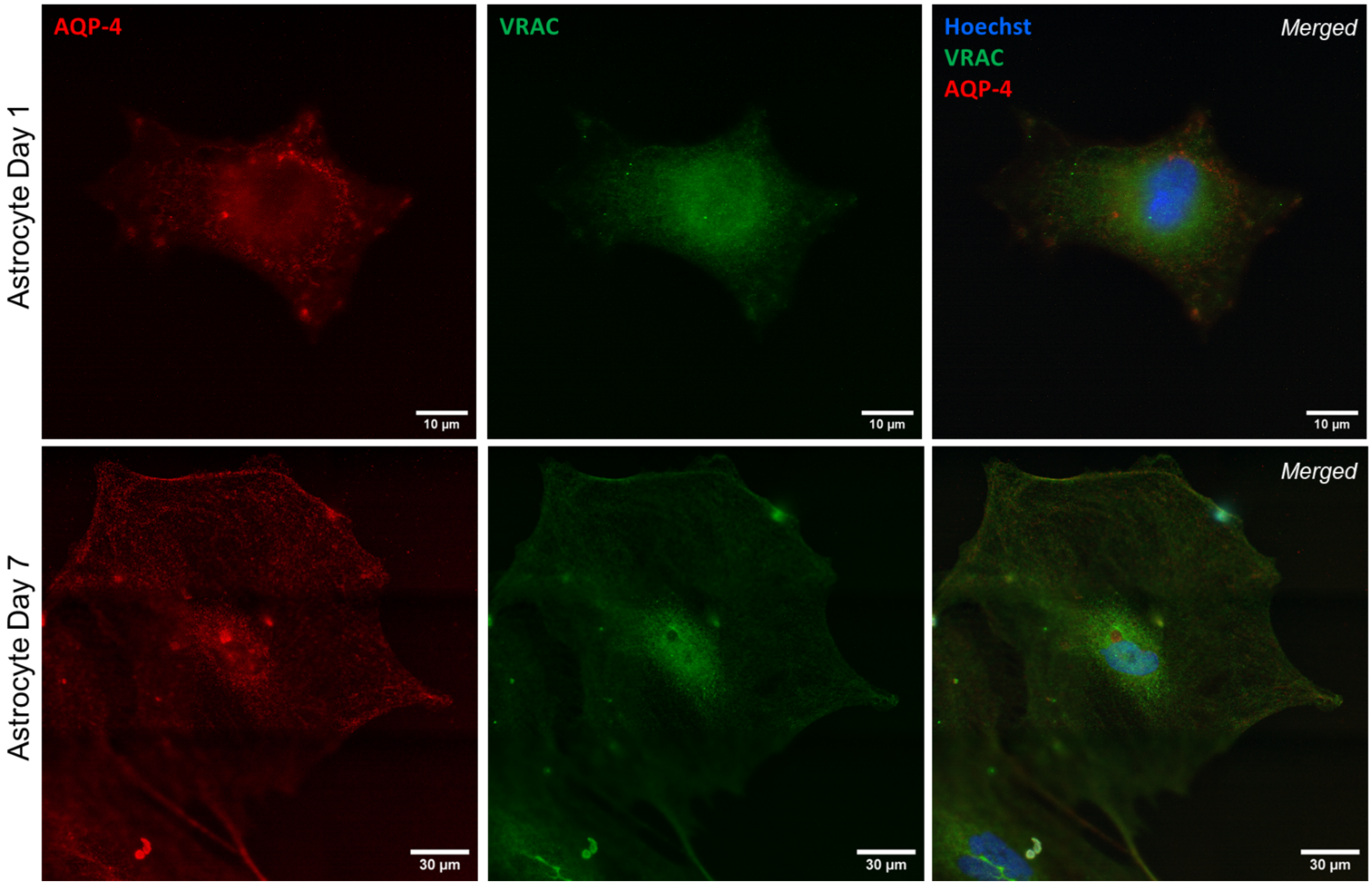


**Figure S1:** QPM images of aquaporin-4 (AQP-4, red), volume-regulated anion channel (VRAC, green), and nucleus (blue) staining of day 1 (top row) and day 7 (bottom row) rat cortical astrocytes.

**Table S1**: P-values and effect size values for various groups of various morphological parameters of postnatal day 2 rat cortical astrocytes.

| Mass | | | |
|---|---|---|---|
| Groups | | P-value | Effect size value |
| Day 1 | Day 3 | 0.5464 | 0.0214 |
| Day 1 | Day 5 | 2.22E-05 | 0.1552 |
| Day 1 | Day 7 | 9.17E-04 | 0.1134 |
| Day 3 | Day 5 | 4.11E-06 | 0.1755 |
| Day 3 | Day 7 | 0.0105 | 0.0917 |
| Day 5 | Day 7 | 8.55E-13 | 0.264 |

| Volume | | | |
|---|---|---|---|
| Groups | | P-value | Effect size value |
| Day 1 | Day 3 | 1.76E-19 | 0.3205 |
| Day 1 | Day 5 | 6.97E-32 | 0.4301 |
| Day 1 | Day 7 | 5.19E-74 | 0.6228 |
| Day 3 | Day 5 | 3.80E-03 | 0.1102 |
| Day 3 | Day 7 | 2.43E-24 | 0.3647 |
| Day 5 | Day 7 | 8.37E-14 | 0.2755 |

| Density | | | |
|---|---|---|---|
| Groups | | P-value | Effect size value |
| Day 1 | Day 3 | 1.60E-33 | 0.4284 |
| Day 1 | Day 5 | 2.54E-106 | 0.8016 |
| Day 1 | Day 7 | 1.86E-124 | 0.812 |
| Day 3 | Day 5 | 3.30E-41 | 0.5123 |
| Day 3 | Day 7 | 8.75E-52 | 0.5424 |
| Day 5 | Day 7 | 4.57E-02 | 0.0737 |

| Area | | | |
|---|---|---|---|
| Groups | | P-value | Effect size value |
| Day 1 | Day 3 | 8.34E-36 | 0.442 |
| Day 1 | Day 5 | 2.65E-100 | 0.7776 |
| Day 1 | Day 7 | 4.83E-144 | 0.8734 |
| Day 3 | Day 5 | 1.41E-32 | 0.4518 |
| Day 3 | Day 7 | 4.90E-80 | 0.6765 |
| Day 5 | Day 7 | 4.93E-22 | 0.356 |

| Area to Volume Ratio | | | |
|---|---|---|---|
| Groups | | P-value | Effect size value |
| Day 1 | Day 3 | 8.34E-07 | 0.1743 |
| Day 1 | Day 5 | 6.50E-54 | 0.5658 |
| Day 1 | Day 7 | 9.02E-62 | 0.5668 |
| Day 3 | Day 5 | 1.27E-30 | 0.4376 |
| Day 3 | Day 7 | 3.85E-38 | 0.4611 |
| Day 5 | Day 7 | 4.00E-03 | 0.1064 |

| Thickness | | | |
|---|---|---|---|
| Groups | | P-value | Effect size value |
| Day 1 | Day 3 | 1.79E-04 | 0.113 |
| Day 1 | Day 5 | 1.88E-20 | 0.3392 |
| Day 1 | Day 7 | 2.27E-24 | 0.3486 |
| Day 3 | Day 5 | 1.81E-11 | 0.256 |
| Day 3 | Day 7 | 3.69E-14 | 0.2712 |
| Day 5 | Day 7 | 1.47E-01 | 0.0536 |
